# Dissecting Molecular Origins of the Mechano-Adaptive Behaviors of Actomyosin Bundles

**DOI:** 10.1101/2025.07.09.663940

**Authors:** Shihang Ding, Tsubasa S. Matsui, Roland R. Kaunas, Taeyoon Kim, Shinji Deguchi

## Abstract

Actomyosin bundles, composed mainly of actin filaments, myosin II filaments, and actin cross-linking proteins, are central to force generation and maintenance in non-striated muscle cells. Actomyosin bundles exhibit diverse mechano-adaptive responses to external mechanical cues. However, the molecular mechanisms underlying such adaptability remain poorly understood. In this study, we employed experiments and agent-based simulations to illuminate how actomyosin bundles adapt to complex mechanical perturbations. Our experiments revealed that individual stress fibers isolated from cells, and thus operating independently of cellular regulation, exhibited buckling in response to rapid compressions following stretch, accompanied by gradual recovery to a straight configuration. Such buckling was not observed under slow compression, indicating that the bundles exhibit rate-dependent adaptive responses. These findings demonstrate that mechano-adaptive behavior is an intrinsic property of actomyosin bundles. Simulations successfully reproduced these rate-dependent behaviors intrinsic to actomyosin bundles and identified key factors determining the dependence. Bundle stiffness influenced the buckling-induced curvature but had little effect on the time required for recovery to straight configuration. Motor density did not alter the recovery time but significantly influenced the magnitude of tensile forces developed after full recovery. The recovery time was primarily governed by the motor walking rate and the applied compressive strain rate. This work reveals how intrinsic mechanics and motor activity drive actomyosin bundle behavior in cellular mechano-adaptation.

## INTRODUCTION

Mechanical forces play an essential role in various biological processes including morphogenesis, migration, and cytokinesis [1-4]. Striated muscle cells in mammals generate contractile forces via molecular interactions occurring in periodic functional units called sarcomeres [5,6]. The sarcomere is composed mainly of actin filaments (F-actin), myosin II, and actin cross-linking proteins (ACPs) such as α-actinin [7,8]. In each sarcomere, the barbed ends of actin filaments are attached symmetrically to Z-disks located on both ends, and thick filaments formed by skeletal muscle myosin II are located at the center of the sarcomere in the M-line. This structure is considered ideal for force generation and contraction.

Non-striated muscle cells that include smooth muscle cells also have contractile machinery consisting mainly of actin filaments, myosin II, and ACPs, but its structure is more disorganized than that of sarcomeres. Actomyosin bundles called stress fibers (SFs) are considered a major source of contractile forces that regulate diverse cellular functions in non-striated muscle cells [11,12]. SFs are categorized into three types, depending on their location and orientation: ventral SFs, dorsal SFs, and transverse arcs [13]. While dorsal SFs are non-contractile due to a very low number of myosins, ventral SFs and transverse arcs are contractile, showing the complementary distributions of domains enriched either with myosin filaments or α-actinin [14]. Structural similarities between sarcomeres and SFs have been discussed in terms of the repeated myosin-enriched domain [15,16], but there are fundamental differences in their structural organization. First, SFs do not have a structure like the M-line found in sarcomeres that can constraint myosin thick filaments near the center.

In addition, the repeated domains with myosin filaments and α-actinin are not uniform in size, and the number of the domains in each SF highly varies, depending on local tension [23,24]. These structural differences might be attributed to their different biological roles. Striated muscles are optimized for unidirectional contraction and force generation, whereas non-striated muscle cells are exposed to more complex mechanical cues exerting in multiple directions. For example, vascular cells, including endothelial cells and smooth muscle cells, are subjected to cyclic stretch with time-varying magnitude and frequency [17-19]. SFs in non-striated muscle cells are not solely specialized for force generation but can inherently adapt to accommodate dynamic mechanical environments. Upon the loss of this adaptability, cells undergo pro-inflammatory signal activation, eventually leading to chronic inflammation-related diseases such as atherosclerosis [20,21]. Given their critical roles, the responses of SFs to mechanical cues have been extensively studied. For instance, when SFs were subjected to small compressive strain (16%) in vivo, the repeated domains in SFs were not compressed equally [25], whereas significant buckling of SFs was observed when they were subjected to larger compressive strain (40%) [26]. In addition, high extensibility of SFs was observed in previous experiments in vitro, which cannot be explained by the conventional models of striated muscle sarcomeres [27].

Although the major components of SFs have been well characterized [28,29], the molecular origins of the mechanical behaviors of SFs remain less clear. Several theoretical and numerical models have been developed for understanding the molecular mechanisms. For example, SFs were simplified as a combination of elements such as springs, dampers, and contractile force generators [30,31], and this assumption was extended to the cellular scale by applying it to finite element models [32-34]. These continuum-based models were able to recapitulate the basic mechanical properties of SFs, but they failed to capture molecular interactions between cytoskeletal proteins. In this regard, agent-based models, which consider each component a separate “agent” and interactions between the agents, have been employed for simulating SFs. Previous agent-based models have provided insights into understanding the role of each molecular component in the mechanical behaviors of SFs, but those models had critical limitations including the oversimplification of myosin filaments and ACPs [35-42].

In this study, using a combination of experiments and agent-based simulations, we investigated the mechano-adaptive behaviors of actomyosin bundles. Our experiments performed with individual SFs isolated from cells demonstrate the rate-dependent behaviors of the SFs. Our model, which rigorously represents the structural and dynamic properties of cytoskeletal components within SFs [43-48], reproduced the experimental observations and also identified critical factors governing the morphological and dynamic behaviors of actomyosin bundles. We also found the molecular mechanism of the mechano-adaptive behaviors of the bundles.

## METHODS

### Experimental characterization

#### Cell culture

Bovine aortic dedifferentiated smooth muscle cells (B354-05, Cell Application), with a proliferative phenotype [49], are cultured with low-glucose (1.0 g/L) Dulbecco’s Modified Eagle Medium (Invitrogen) containing 10% (v/v) heat-inactivated fetal bovine serum (SAFC Biosciences) and 1% penicillin–streptomycin (Invitrogen) in a 5% CO_2_ incubator at 37°C.

#### Isolation and manipulation of individual SFs

To isolate individual SFs from cells cultured on a glass-bottom dish (Iwaki), we first remove the apical cell membrane, cytosol, and nuclei via the hypotonic shock technique followed by a gentle detergent treatment using the method described elsewhere (Fig. 1A) [29,50,51]. The resulting isolated SFs are immersed within the activation solution prepared as follows: 2 mM free Mg^2+^, 20 mM imidazole, 2.1 mM CaCl_2_, 2 mM EGTA, and 1 mM Mg-ATP. It is observed that the isolated SFs still maintain contractility in similar Mg-ATP buffer (Fig. S1A). One tip of the remaining ventral SFs is isolated from the underlying substrate by using a glass needle. The glass needle is manufactured using a glass-electrode puller (P-1000, Sutter Instrument) to have sufficient stiffness to resist bending induced by a force generated by SFs. The glass needle is functionalized in advance with 2% (3-aminopropyl) trimethoxysilane (Sigma-Aldrich) in 90% EtOH for 30 min and subsequently with 20% glutaraldehyde (Wako) for 5 min. The lifted individual SF, with the other end remaining attached to the substrate, is stretched and then compressed using the glass needle (Figs. 1B and S1B). The response of SFs is observed under phase-contrast microscopy (IX-71, Olympus) imaged with a CCD camera (ORCA-R2, Hamamatsu) at room temperature.

**Fig. 1.**
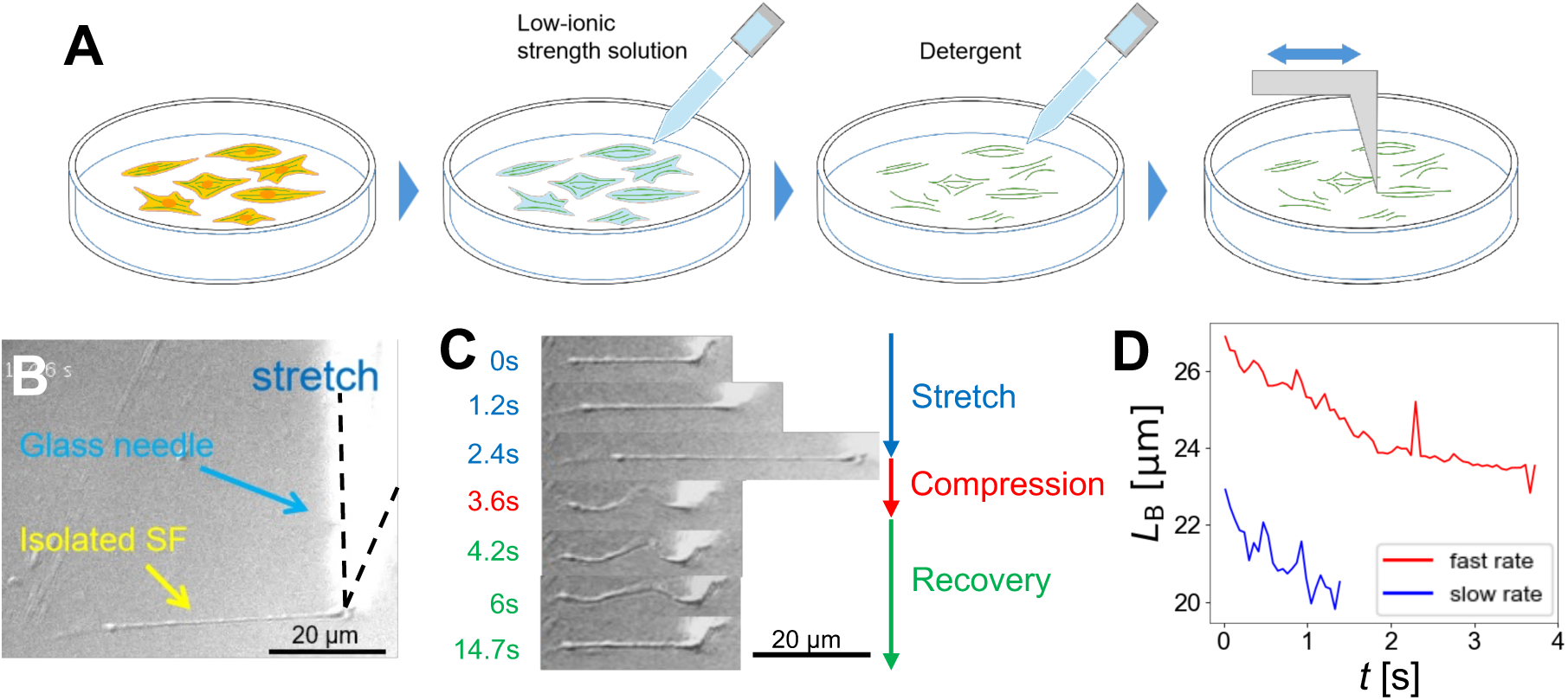
Characterization of the mechanical behaviors of a stress fiber (SF) isolated from cells. (A) A single SF was isolated from A7r5 cells by removing an apical cell membrane, cytosol, and a nucleus via the hypotonic shock technique. The SF remained attached to an underlying substrate by treating the de-roofed cells with a detergent. Further manipulation was conducted using a functionalized glass needle. (B) An individual SF was stretched by the glass needle (black dashed lines). (C) Manipulation of the single SF in stretch (blue), compression (red), and recovery (green) phases. Bucking of the SF was often observed after the compression phase. During the recovery phase, the SF gradually became straight. (D) The contour length (*L*_B_) of SFs during the recovery phase over time. Faster compression (red) resulted in longer *L*_B_, whereas slow compression (blue) led to shorter *L*_B_. However, the rate of a decrease in *L*_B_ was similar between two cases.

### Agent-based model

#### Model overview and simulation setup

In our model, cytoskeletal components – actin filaments, ACPs, and motors – are simplified using cylindrical segments connected by elastic hinges (Fig. S2A). ACPs transiently bind to actin filaments to form cross-linking points. Motors mimicking the structure of myosin filaments bind to and walk along actin filaments. The positions of the cytoskeletal components are updated every time step using the Langevin equation and the Euler integration scheme. Details about the mechanics and dynamics of the components and all parameters used in our model are explained in Supplementary Text and Supplementary Table.

At the beginning of simulations, *N*_F_ filaments whose length is 4.9 μm are allocated in a rectangular computational domain (20×20×5 μm) (Fig. 2A). The initial orientation of all the filaments is in the z direction. The barbed ends of the half of the filaments are connected to the -z boundary (z = 0) via a spring whose stiffness is *κ*_s,bnd_, and the barbed ends of the other half are connected to the +z boundary (z = 5 μm) via the same type of spring. This configuration mimics the structure of one contractile unit found in muscle cells and SFs. The x and y positions of these filaments are located along a circle whose diameter is 27 nm. After the allocation of actin filaments in the domain at the beginning of simulations, ACPs and motors are allowed to bind to actin filaments. During this step, the actin filaments are not allowed to move, and motor arms do not walk. Admittedly, it is not possible to allocate more than 12 filaments (whose diameter is 7 nm) along this circle without physical overlaps. Nevertheless, we chose this way to let all filaments be cross-linked to each other by a similar chance for more uniform connectivity between filaments; if a large number of filaments are allocated without any physical overlap, some of the filaments cannot be interconnected because a distance between them can be higher than what ACPs or motors can reach. As a result of this assembly process, a bundle is created with two sets of filaments with opposite polarity, interconnected by motors and ACPs. Unless specified, the molar ratios of motors (*R*_M_ = *C*_M_ / *C*_A_) and ACPs (*R*_ACP_ = *C*_ACP_ / *C*_A_) are set to 0.01 and 0.1, respectively, where *C*_A_, *C*_M_, and *C*_ACP_ indicate the concentrations of actin, motors, and ACPs, respectively.

**Fig. 2.**
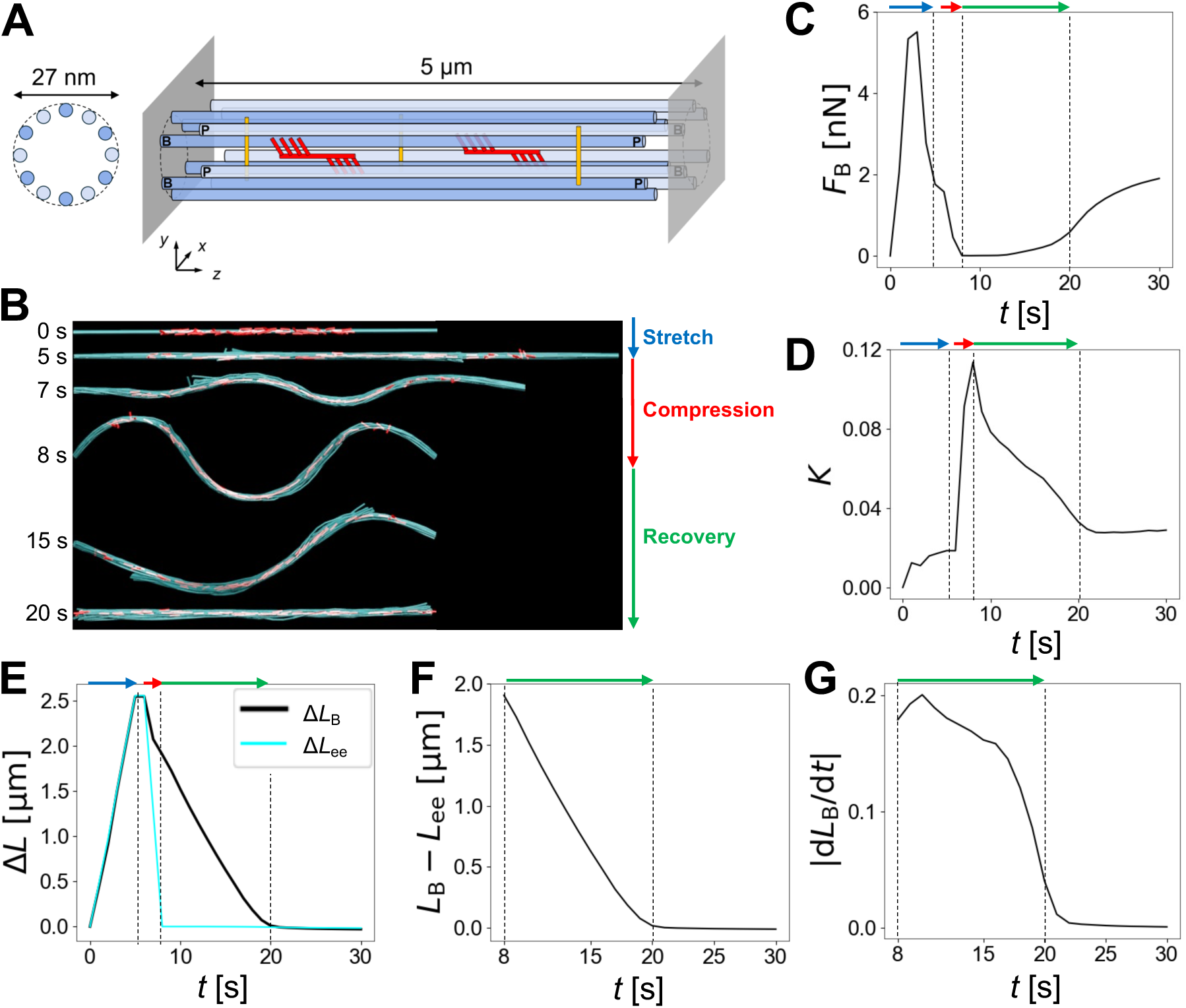
An actomyosin bundle simulated via the agent-based model reproduced the experimental observations. (A) In our model, a bundle consists of two sets of actin filaments with opposite polarities (dark blue and light blue). “B” and “P” indicate the barbed and pointed ends of the filaments, respectively. Their length is 4.9 μm which is close to the initial length of the bundle, 5 μm. The barbed ends of actin filaments are connected to either the left or the right boundary via springs. Motors (red) and actin cross-linking proteins (ACPs, yellow) are added to the bundle. (B) An example of simulations for testing the stretch, compression, and recovery of the bundle. Buckling emerged during the compression, followed by gradual recovery into a straight shape, which is consistent with the experiment results. Only actin filaments (cyan) and motors (red) are visualized in these snapshots. (C) Average tensile force acting on the bundle (*F*_B_) over time. A large force is developed and then partially relaxed during the stretch phase. The force is fully relaxed during the compression phase, and it increases again after the bundle becomes fully straight. (D) The curvature of the bundle (*K*) over time. A sharp increase in *K* was observed during the compression phase, followed by a graduate decrease in the recovery phase. (E) A comparison between a change in bundle contour length (Δ*L*_B_, black) and a change in the end-to-end distance (Δ*L*_ee_, cyan). (F) A difference between *L*_B_ and *L*_ee_ over time. (G) The magnitude of the decreasing rate of *L*_B_ over time. In (C-G), the stretch, compression, and recovery phases are indicated by arrows with the same colors as those used in (B). Vertical dashed lines indicate the end of each phase. Note that there is time delay of 1 s between the end of the stretch phase and the beginning of the compression phase.

#### Stretch, compression, and recovery tests

After the completion of bundle formation, actin filaments are allowed to move, and motor arms are allowed to walk along the actin filaments. Due to the repulsive forces acting between filaments and the thermal fluctuation of filaments, the bundle becomes thicker with the spatial redistribution of filaments to some extent. Then, the +z boundary where the barbed ends of the half of filaments are attached is displaced in the +z direction, following a linearly increasing strain. The stretching rate during this phase is *ε̇*_s_ = 0.1 s^-1^ (Fig. 2B). After reaching the maximum strain level (*ε* = 0.5), the boundary stays at the maximum strain level for 1 s for further force relaxation. Then, the boundary is displaced in the -z direction with the compressive rate (*ε̇*_c_) until it returns to the initial position, *ε* = 0. This is called the compression phase. Then, the strain is held at 0, and the behaviors and force of the bundle are quantified.

#### Quantification of bundle force, curvature, and contour length

To quantitatively measure a tensile force with high temporal resolution, the domain is divided into 20 cross-sections equispaced in the z direction. All segments that pass each cross-section are selected, including actin segments, motor arms, motor backbone segments, and ACP segments. Extensional forces acting on those segments are measured, and the z component of those forces is summed at each cross-section. Then, 20 sums are averaged to obtain a single quantity (*F*_B_) representing a tensile force acting on the entire bundle (Fig. 2C). This calculation is done every 1 s.

The curvature of the bundle is calculated using the three-dimensional positions of actin segments. First, at each endpoint of filament segments, the first and second derivative vectors are calculated using vectors defined by two endpoints of segments adjacent to the point. Local curvature at the point is then determined by calculating cross product between the derivative vectors:

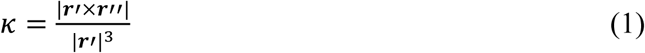

where *κ* is the local curvature at each endpoint, ***r***′ and ***r***′′ are the first and second derivative vectors, respectively. The norm of the cross product is divided by the cube of the first derivative to obtain the local curvature at each endpoint. The bundle-level curvature is obtained by averaging the mean curvature of all the actin filaments (Fig. 2D).

The end-to-end distance of the bundle (*L*_ee_) is calculated by measuring a distance between the leftmost and rightmost endpoints of actin segments in the z direction. To quantify the contour length of the entire bundle (*L*_B_), two contour lengths are separately measured in two half-domains divided in the z direction (Fig. S2B). For calculating each of the two contour lengths, only one set of filaments (whose barbed ends are attached to a boundary on that side) is used. The length of each filament in the set, which belongs to the half-domain, is calculated by summing the length of actin segments located within the half-domain. This filament length is averaged over all filaments in the set for the half-domain. Then, these two contour lengths are added into *L*_B_ (Fig. 2E).

## RESULTS

### SFs exhibit rate-dependent responses and mechano-adaptability

In our experiments, we stretched and then compressed individual SFs isolated from cells, via a functionalized glass needle at different rates. SFs were buckled after compression and then gradually recovered into a straight shape (Figs. 1C and S1). We found that the SFs compressed with a higher strain rate showed more significant buckling and required longer time to recover into the straight shape (Fig. 1D; see Supplementary Video S1 for detail). Interestingly, the rate of a decrease in the bundle length was similar regardless of the compression rate. These observations suggest that SFs exhibit rate-dependent responses and possess the ability to mechanically adapt to external cues by elongating, buckling, and shortening across a range of strain rates. We confirmed that the primary constituents of these isolated SFs are identical to those of intracellular SFs [28,29]. Therefore, the only biochemical difference between intracellular and extracellular SFs is whether or not the turnover of protein constituents via molecular exchange with their surrounding takes place. Our results demonstrate that, even without such protein turnover, SFs can undergo repeated stretching and contraction.

At first glance, these behaviors of SFs may appear to arise from the viscoelastic retraction of SFs proposed in a previous study [52]. However, viscoelastic models consisting of dashpot(s) and spring(s), which are commonly used for fitting the viscoelastic responses of materials, cannot explain these behaviors [53]. For example, the Kelvin-Voigt model with one spring and one dashpot connected in parallel could explain the gradual SF retraction after laser ablation [52], but this model cannot be stretched abruptly because of the nature of the dashpot. In addition, the Maxwell model with one spring and one dashpot connected in series can be stretched instantaneously, but it cannot explain the gradual recovery. The standard linear solid model with two springs and one dashpot cannot explain the gradual recovery either, suggesting that our observations are not directly related to the viscoelastic nature of SFs. We hypothesized that the mechanical response of SFs to perturbations may involve force-dependent unbinding and walking of myosin II.

### A minimal bundle reproduced buckling and gradual recovery

To verify our hypothesis and find factors inducing buckling and gradual recovery, we employed our well-established agent-based computational model (Fig. 2A). First, we tried to reproduce our experimental results showing buckling and gradual recovery after compression. A minimal bundle in the model consists of two sets of filaments with opposite polarity connected by motors and ACPs, which mimics the structure of a single contractile unit existing in sarcomeres or SFs. When the bundle was stretched up to the strain of *ε* = 0.5 for 5 s (i.e., a tensile strain rate, *ε̇*_s_= 0.1), a force acting on the bundle (*F*_B_) highly increased at the beginning, but *F*_B_ was significantly reduced before the end of the stretch phase (Figs. 2B, C). This force relaxation was attributed to the loss of connections between the two sets of filaments. Most of the motor arms and ACPs, which inter-connected actin filaments in the anti-parallel configuration, dissociated from actin filaments due to high tensile forces developed on them. Despite the possibility of rebinding to actin filaments after unbinding, tensile forces that filaments previously bore were relaxed after each dissociation event, resulting in a permanent increase in the length of the bundle with a shorter overlap between the two sets of filaments and the reduction of *F*_B_. The strain was maintained at 0.5 for 1 s (pausing), and then the bundle was compressed back to *ε* = 0 for 2 s (i.e., a compressive strain rate, *ε̇*_c_= -0.25). During the compression phase, the curvature of the bundle (*K*) rapidly increased, followed by a gradual decrease (Fig. 2D). The contour length of the bundle (*L*_B_) was similar to the end-to-end distance (*L*_ee_) during the stretch and pausing phases. Even after force relaxation, there is a residual tensile force at the beginning of the compression phase (Fig. S3), inducing elastic recovery with a small, abrupt drop in *L*_B_. Then, *L*_B_ decreased slower than *L*_ee_ during the compression phase (Fig. 2E), resulting in the buckling. A difference between *L*_B_ and *L*_ee_ became maximal at the end of the compression phase (Fig. 2F). After compression, *L*_B_ initially decreased with a relatively constant rate, but the decreasing rate was reduced as *L*_B_ approached *L*_ee_ (Fig. 2G). When the difference between *L*_B_ and *L*_ee_ becomes smaller than 0.01 μm, the bundle is considered to reach the end of the recovery phase. *F*_B_ started increasing near the end of the recovery phase because motors can build up tensile forces better on filaments in a straight bundle by feeling resistances from boundaries where the barbed ends of filaments are attached (Fig. 2C).

### Bundle stiffness affects a buckling pattern but has a minimal effect on recovery time

We have showed the mechanical and dynamic behaviors of the bundle at the stretch, compression, and recovery phases. To identify determining factors for each behavior, we performed parametric studies by changing one parameter with the others fixed.

First, we varied bundle stiffness by changing the bending stiffness of actin filaments (*κ*_b,A_). Note that we used 10-fold higher bending stiffness for actin filaments under the reference condition than an experimentally measured value 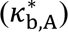 to achieve a clear buckling pattern 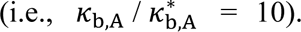 *κ*_b,A_ was reduced to 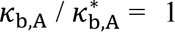 or increased to 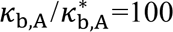 [54], [55]. The direct impact of the variation in *κ*_b,A_ was observed in *K* measured at the end of the compression phase (*K*_r_) (Figs. 3A, B); with higher *κ*_b,A_, *K*_r_ was noticeably lower. *L*_B_ measured at the end of compression and time required for full recovery (*t*_r_) were almost independent of *κ*_b,A_ (Figs. 3C, D). Time evolution of *L*_B_ and *F*_B_ were similar in cases with different *κ*_b,A_ (Fig. S4). These results imply that the mechano-adaptive behavior of the bundle was not regulated by the bundle stiffness.

**Fig. 3.**
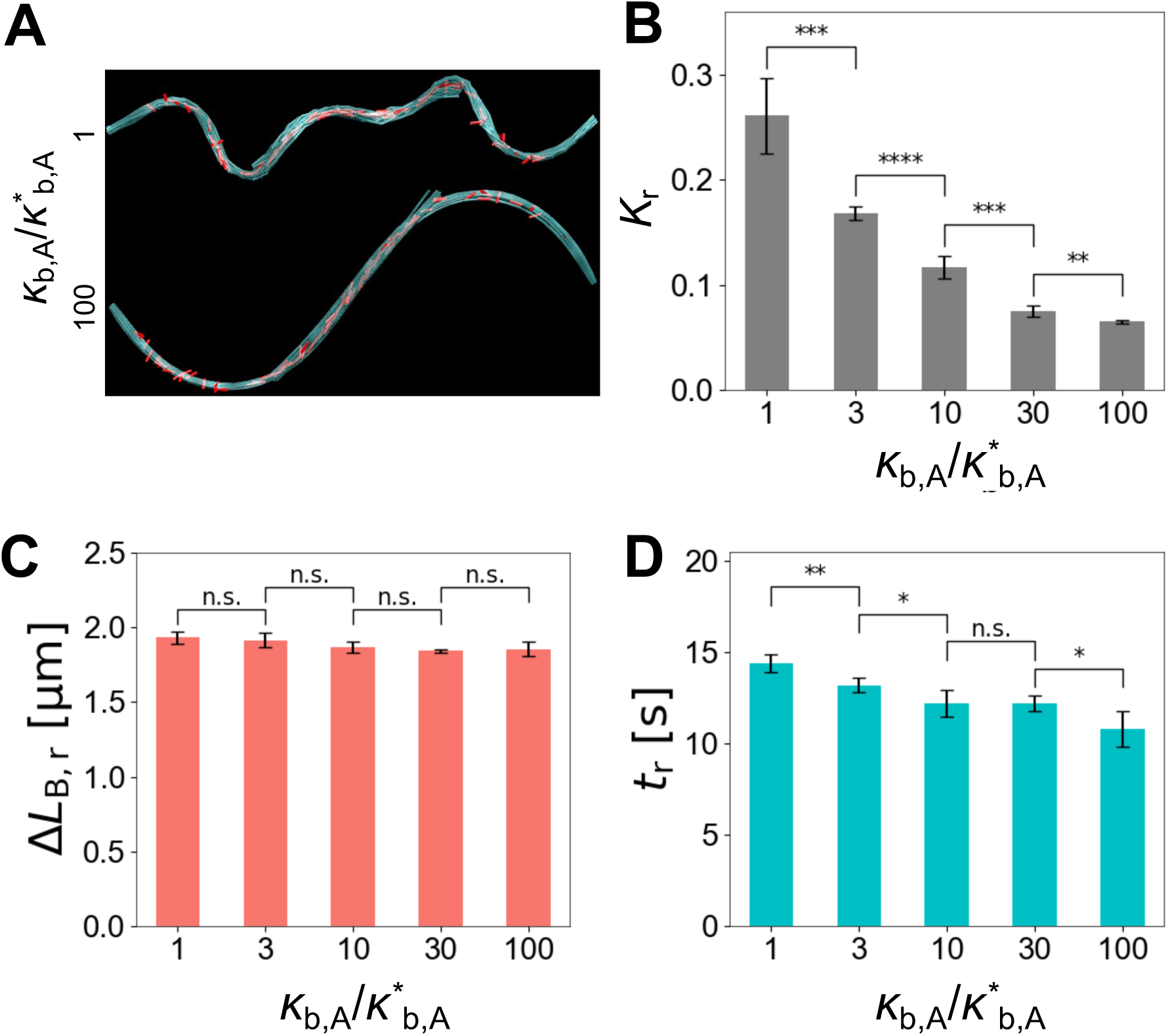
Bending stiffness of actin filaments (*κ*_b,A_) affected the bundle curvature but hardly varied recovery time. (A) The shape of bundles at the end of the compression phase with different *κ*_b,A_. *κ*^∗^ represents the reference value of *κ*_b,A_. Only actin filaments (cyan) and motors (red) are visualized in these snapshots. (B-D) Bundle curvature (*K*_r_) and a change in bundle contour length (Δ*L*_B,r_) measured at the end of the compression phase and time taken for full recovery into a straight shape, *t*_r_, with different *κ*_b,A_. With higher *κ*_b,A_, bundles showed smaller curvature despite similar bundle contour length, but it took similar time for the bundle to fully recover.

### How fast the bundle is compressed significantly affects buckling and recovery pattern

Motivated by our experimental observations (Fig. 1D), we varied *ε̇*_c_ which determines how fast the bundle is compressed after stretching. As a bundle was compressed faster, the bundle showed higher contour length and larger curvature at the end of compression, and it took longer time for the bundle to recover into a straight shape (Figs. 4A-D), which is consistent with experimental observations. (i.e., *L*_B_, *K*_r_, and *t*_r_ were proportional to |*ε̇*_c_|.) Note that different *K*_r_ in these cases is related to a difference in *L*_B_, not filament bending stiffness, *κ*_b,A_; to fit a bundle with longer contour length to the same end-to-end distance, the curvature of the bundle should be larger. With the smallest |*ε̇*_c_|, *L*_B_ was close to *L*_ee_, so buckling was not observed (Figs. 4A, top and S5A, B). In addition, *F*_B_ was much larger during the compression phase than that in other cases because a relatively straight bundle during compression in this case allowed motors to generate large tensile forces (Fig. S5D). Except this case, the recovery rate in other cases was similar to each other (Fig. S5C), implying that the bundle shortening rate was not determined by |*ε̇*_c_|. We hypothesized that the dependence of the bundle behaviors on |*ε̇*_c_| is attributed to the walking of motors during the compression phase. As it takes more time for compressing the bundle, motors can walk over a longer distance on two sets of filaments with opposite polarity, which results in an increase in the overlap between the filaments and therefore leads to a decrease in *L*_B_ at the end of compression.

**Fig. 4.**
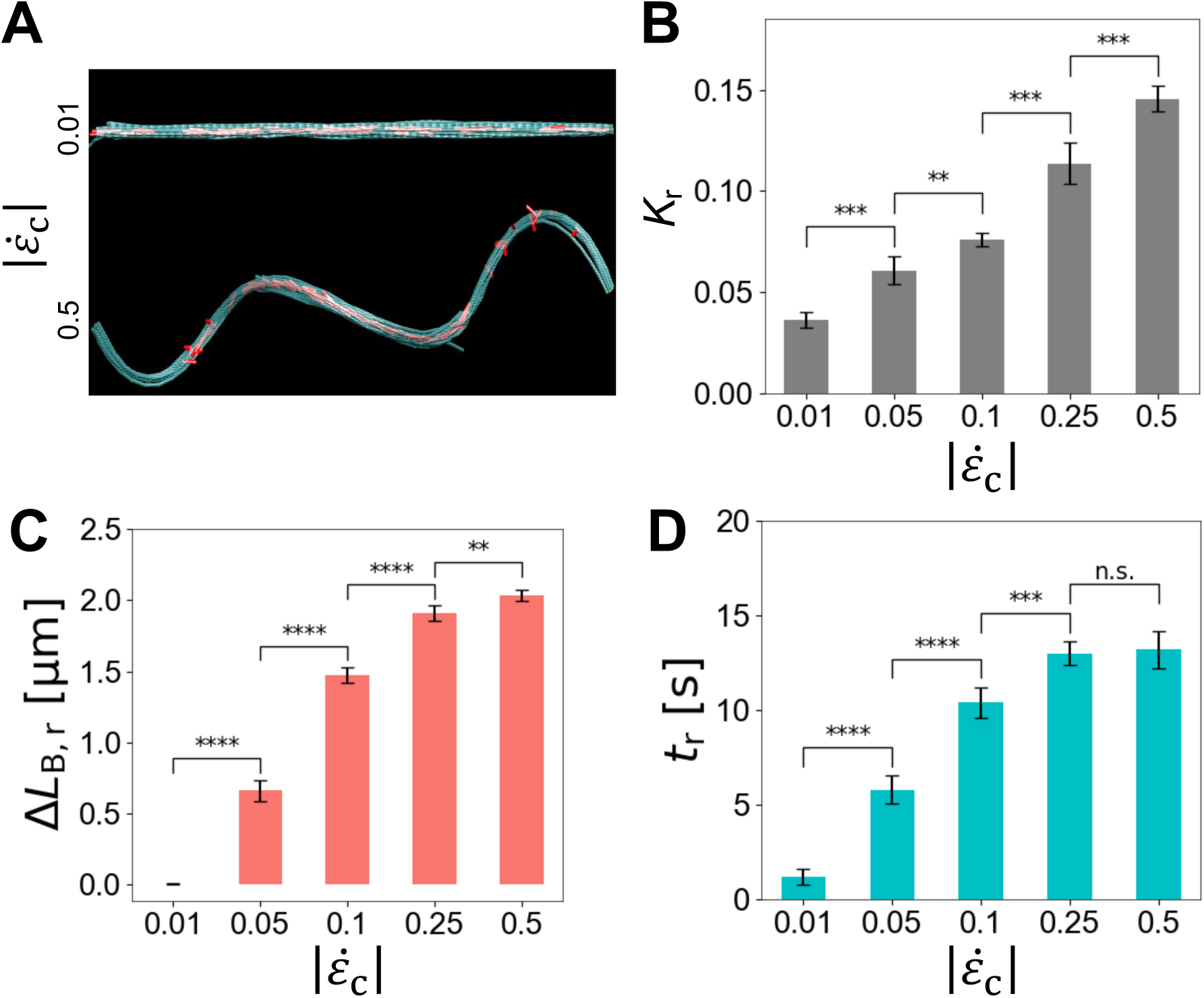
Slowly compressed bundles did not exhibit significant buckling. (A) The shape of bundles right after full compression with the different magnitude of compression rates (|*ε̇*_c_|). Only actin filaments (cyan) and motors (red) are visualized in these snapshots. (B-D) Bundle curvature (*K*_r_) and a change in bundle contour length (Δ*L*_B,r_) measured at the end of the compression phase and time taken for full recovery into a straight shape, *t*_r_, depending on |*ε̇*_c_|. As bundles were compressed slower, the contour length was shorter after full compression, resulting in lower curvature and shorter recovery time.

### Motor walking speed directly impacts the recovery phase, not motor density

If our hypothesis is correct, a change in the walking speed of motors would impact the contour length and recovery time of the bundle. To test this, we varied the walking rate of motors by varying one of the mechanochemical rates used in the parallel cluster model that determines the unbinding and walking rates of motors. This rate is called the ATP-dependent unbinding rate (*k*_20_) which defines a transition rate from the post-power-stroke state to the unbound state. With higher *k*_20_, the walking and unbinding rates of motors are enhanced (Fig. S6). With higher *k*_20_, *L*_B_ measured at the end of the compression phase was smaller because motors walked over a longer distance during the bundle compression (Figs. 5A and S7C, D). However, since the duration of the compression phase was only 2 s, this difference in *L*_B_ originating from different walking speed cannot be very large in these cases. As a result, *K*_r_ was slightly smaller with higher *k*_20_ (Figs. S7A, B). It took much shorter time for the bundle to recover with higher *k*_20_. This is partially attributed to smaller *L*_B_ at the beginning of the compression phase, but we found that the decreasing rate of *L*_B_ (i.e., the bundle shortening rate) was proportional to *k*_20_ (Figs. 5C and S7E), corroborating our hypothesis that a decrease in *L*_B_ during the recovery phase is driven mainly by the walking of motors. *F*_B_ was independent of *k*_20_ until the end of the recovery phase, but *F*_B_ increased faster with higher *k*_20_ after full recovery into a straight shape (Fig. S7F).

**Fig. 5.**
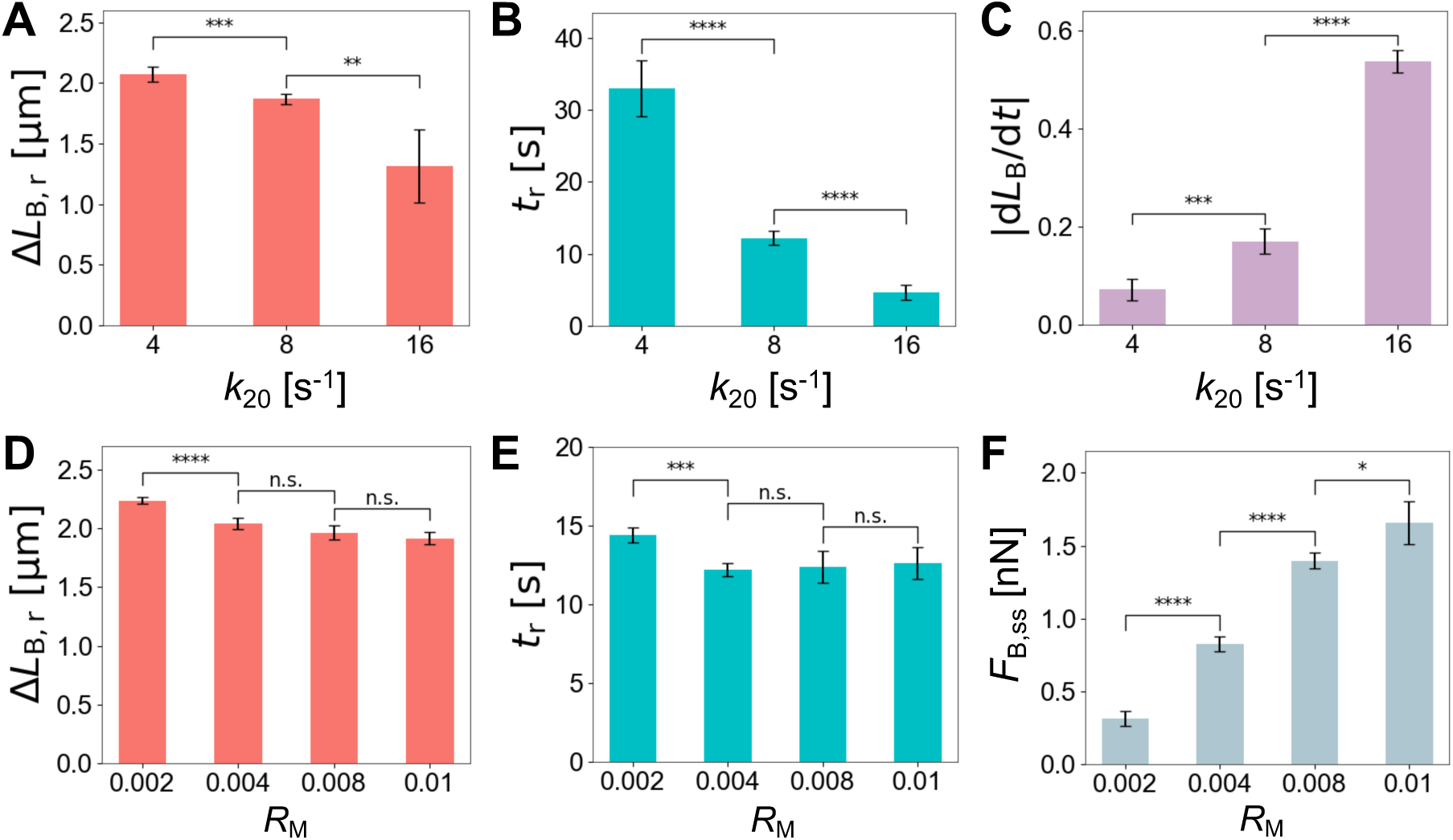
The recovery rate of the bundle is determined by motor walking speed, not by motor density. (A-C) *k*_20_ is the ATP-dependent unbinding rate of myosin heads used in the parallel cluster model, corresponding to a transition rate from the post-power-stroke state to the unbound state. In general, higher *k*_20_ for motors results in higher unbinding and walking rates. (A, B) A change in bundle contour length (Δ*L*_B,r_) at the end of compression and time taken for full recovery (*t*_r_) with different *k*_20_. (C) The decreasing rate of *L*_B_ right after the end of compression. As motors walk faster, it took much less time for the bundle to fully recover. (D, E) Δ*L*_B,r_ and *t*_r_ with different motor density (*R*_M_). (F) Average tensile force acting on the bundle at the steady state after full recovery (*F*_B,ss_), depending on *R*_M_. With more motors, a larger force was generated after full recovery, but the recovery time hardly changed.

Next, we probed the effects of motor density (*R*_M_) (i.e., the molar ratio of motors) on the dynamic and mechanical behaviors of the bundle. When we reduced *R*_M_ from the reference value 0.01, we did not notice any meaningful change in the contour length and curvature of the bundle at the end of compression and recovery time (i.e., *L*_B_, *K*_r_, and *t*_r_ were largely independent of *R*_M_.) (Figs. 5D, E and S8A-E). The only difference was the level of *F*_B_ after the end of full recovery; with higher *R*_M_, *F*_B_ became higher (Figs. 5F and S8F). These results imply that the number of motors in the bundle does not mediate bundle behaviors during the stretch, compression, and recovery phases.

## DISCUSSION

A large fraction of mechanical forces in cells are generated by actomyosin bundles. The adaptive behavior of the actomyosin bundles to mechanical environments is intimately related to diverse biological functions. In this study, we combined our agent-based computational model with experimental findings to delve into the molecular mechanism of the mechano-adaptative behavior of actomyosin bundles. In our experiments, we observed that individual SFs exhibited diverse responses depending on loading conditions; SFs showed buckling with high curvature after rapid compression following stretch, and then the contractile SFs gradually recovered to a straight shape (Figs. 1 and S1), which was not observed with slow compression. These results suggest that the permanent elongation of SFs induced by large stretch, which initially appeared irreversible, was reversed. These observations show the intrinsic mechanical properties of individual SFs that are obscured by the complex interplay of multiple factors within cells. Thus, this is the first experimental observation of the mechano-adaptive behavior of SFs, beyond previously characterized mechanical properties of SFs, such as tensile properties [50] and intrinsic contractile forces [51].

The rate-dependent behaviors of the SF were reproduced by our agent-based computational model (Figs. 2 and S3). A minimal bundle in the model could be stretched up to 50% strain and exhibit distinct buckling after compression, followed by gradual recovery to a straight shape. We found that the buckling pattern is determined by two factors. The first factor was bundle stiffness (Figs. 3 and S4) which can be changed by varying either the number or stiffness of filaments. Despite lack of physiological relevance, we varied the filament stiffness, rather than the total number of filaments; to avoid a substantial increase in the computational cost of simulations; since these filaments in the bundle are very close to each other, the number of neighboring pairs, which we should consider for calculating repulsive forces between filaments and for possible binding between filaments and ACP/motor, substantially increases if the bundle has more filaments. Note that the number of actin filaments per cross-section in our simulations is already higher than that estimated for real SFs, 10-30 filaments [56]. It was observed that higher bundle stiffness resulted in the smaller curvature of the bundle after fast compression. The second determining factor for the buckling pattern was the compression rate of the bundle (Figs. 4 and S5). The compression rate determined the contour length of the bundle at the end of compression. Faster compression resulted in greater contour length, resulting in more significant buckling. By contrast, with very slow compression, buckling was not observed at all after compression. During compression, motors can still walk toward the barbed ends of actin filaments without a load, which can reduce the bundle contour length by increasing overlap between two sets of actin filaments with opposite polarity in the bundle. If the unloaded walking speed of motors is greater than the compression rate of the bundle, the bundle can maintain a straight shape during the compression. If not, the bundle contour length will be still higher than the end-to-end distance of the bundle, leading to buckling.

A high tensile force was developed on the bundle at the first half of stretch, but it was relaxed to less than half of the peak force during the last half of stretch. The force gradually decreased to zero during compression. We found that this residual tensile force does not mediate the gradual recovery of the bundle into the straight shape. When we increased the number of motors (Figs. 5D-F and S8), the residual force during compression became higher, but the decreasing rate of the bundle contour length was not changed noticeably because the maximal walking speed of motors is independent of the motor density. We demonstrated that the walking speed of motors determines how fast the bundle became straight after compression (Figs. 5A-C and S7). These observations imply again that motors walk along filaments without a significant load during the bundle recovery. If motors feel a large resistance when they pull a pair of “curvy” anti-parallel filaments in the opposite directions, their walking speed will be reduced to a different extent depending on motor density. Then, the recovery time or the bundle shortening rate will show dependence on motor density. Results in this study are different from an observation in our previous study where larger motor density resulted in the faster contraction of two-dimensional networks [46]. In the networks, actin filaments were randomly located and oriented, so motors contracted the network against intrinsic elastic resistance; as network contraction proceeds, clustering or aggregation of filaments occurs, and filament density highly increases. In the presence of such an elastic resistance, larger contractile forces generated by more motors could lead to faster network contraction. It is expected that disorganized actomyosin bundles consisting of filaments with random locations and orientations would contract faster with more motors.

After the full recovery of the bundle, motors generated large tensile forces on the straight bundle by pulling filaments against a resistance from springs connected to the rigid boundaries. Then, motors walked more slowly due to the developed force and eventually stopped walking when they felt a force beyond the stall force level. Thus, the bundle-level tensile force showed a clear plateau. In this state, the plateau level was proportional to motor density, which is consistent with our previous study showing larger force generation with motor motors on a straight bundle [43].

There was a foundational study showing the gradual retraction of SFs after laser ablation [52]. The retraction was understood as a result of pre-existing tension existing before the ablation. Although the Kelvin-Voigt model consisting of a spring and a dashpot could fit the retraction curve, we postulate that the retraction was partially driven by the walking activities of myosin II. We showed that the decreasing rate of the bundle contour length was initially high but became lower as the contour length approached the initial bundle length before stretch (Fig. 2E). This is qualitatively similar to the retraction pattern observed in the former study, implying that the partial contribution of motor walking to retraction after laser ablation should be considered as well. There have been more advanced viscoelastic models with spring, dashpot, and active contractility that represent myosin activity [57], but these models were not used for explaining bundle retraction under the load-free condition.

One of the differences between experiments and simulations is the rate of bundle recovery. The SFs used in our experiments reduced its length much faster than the bundle in our simulations. One reason should be different motor walking speed. Considering that A7r5 cells are smooth muscle cells, the walking speed of myosin II in the cells would be ∼200 nm/s [58]. This is higher than the unloaded walking speed of motors used in our model, 56 nm/s, but this cannot solely explain the large difference in the bundle shortening rate between experiments and simulations. It has been observed that SFs have a structure with repeated subunits consisting of localized myosin II and α-actinin [59], and each unit structurally mimics sarcomeres found in muscle cells. If this is the case, SFs can reduce their contour length much faster because a decrease in the contour length occurs in multiple locations simultaneously. Our computational model represents only one of these subunits in SFs, which explains a large discrepancy in the bundle shortening rate and the recovery time between experiments and simulations.

## CONCLUSIONS

In this study, motivated by our experimental observations on the rate-dependent behaviors of SFs, we investigated the origins of the rate dependence, using the agent-based model that mimics a repeated subunit of actomyosin bundles found in SFs. Our model reproduced overall experimental observations and further showed how the shape and recovery of the bundles are determined. The bundle shape was mainly determined by bundle stiffness and how fast the bundle was compressed after stretching. The recovery time was determined by the compression rate and how fast motors walk without a load. These results provide insights into understanding of the mechano-adaptive behavior of actomyosin bundles. We will use a similar model to create a SF-like bundle with multiple contractile units to probe the high extensibility and mechanical behaviors of a whole SF.

## ACKNOWLEDGMENTS

We gratefully acknowledge the support from EMBRIO Institute, contract #2120200, the National Science Foundation (NSF) Biology Integration Institute and the support from JST SPRING (JPMJSP2138 to SDi), JSPS (Japan Society for the Promotion of Science) KAKENHI (24680049 and 21H03796 to SDe).

## SUPPLEMENTAL INFORMATION

### Simplification of cytoskeletal elements

In our model, actin filaments are coarse-grained into serially connected cylindrical segments with polarity defined by barbed and pointed ends (Fig. S2A). Actin cross-linking proteins (ACPs) comprise two segments connected at their center point. Motors mimic the structure of myosin filaments; each motor has a backbone structure with 8 arms (*N*_a_ = 8), and each arm kinetically represents 4 myosin heads (*N*_h_ = 4). Therefore, the total number of myosin heads represented by one motor is 32 (= *N*_a_×*N*_h_). The motor backbone consists of 7 segments with identical length, and a segment located at the center corresponds to the bare zone of myosin filaments. Two motor arms are connected to each endpoint of the backbone segments.

### Brownian dynamics via the Langevin equation

The velocities of the endpoints of all the segments constituting F-actin, motor, and ACP are calculated using the Langevin equation with inertia neglected:

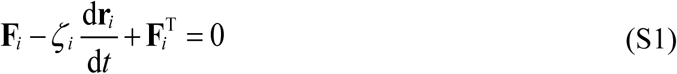

where **r***_i_* is a position vector for the *i*th element, *ζ_i_* is a drag coefficient, *t* is time, **F***_i_* is a deterministic force, and **F**_*i*_^T^ is a stochastic force satisfying the following fluctuation-dissipation theorem [60]:

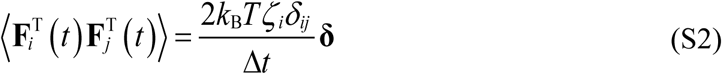

where *δ_ij_* is the Kronecker delta, **δ** is a second-order tensor, *k*_B_*T* is thermal energy, and Δ*t* = 1.15433×10^-6^ s is time step. The drag coefficients are calculated using the following approximated form for a cylindrical object [61]:

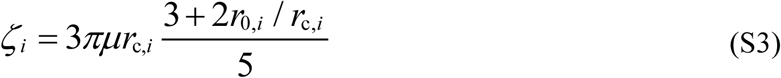

where *μ* is the viscosity of a surrounding medium, and *r*_0,*i*_ and *r*_c,*i*_ are the length and diameter of a segment, respectively. The positions of all the segments are updated each time step via the Euler integration scheme and velocities calculated by Eq. S1 (d**r***_i_*/d*t*):

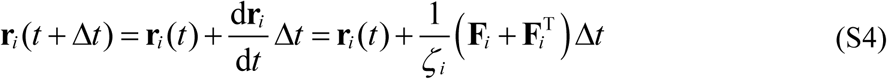

### Structural and mechanical properties of elements

Deterministic forces include extensional forces maintaining equilibrium lengths, bending forces maintaining equilibrium angles, and repulsive forces representing volume-exclusion effects between overlapping actin filaments. The bending and extensional forces originate from the following harmonic potentials:

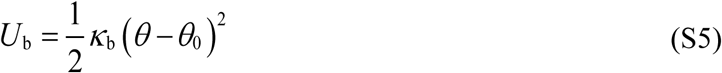

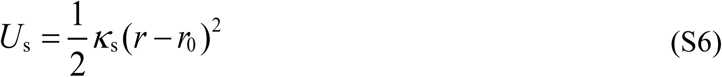

where *κ*_b_ and *κ*_s_ are bending and extensional stiffnesses, *θ* and *θ*_0_ are instantaneous and equilibrium angles formed by adjacent segments, and *r* and *r*_0_ are the instantaneous and equilibrium lengths of cylindrical segments. An equilibrium angle formed by two adjacent actin segments (*θ*_0,A_ = 0 rad) and the equilibrium length of each actin segment (*r*_0,A_ = 140 nm) are maintained by bending (*κ*_b,A_) and extensional (*κ*_s,A_) stiffnesses of actin filaments, respectively. The reference value of *κ*_b,A_ (= *κ*\*_b,A_) corresponds to the persistence length of 9 μm (4). An equilibrium angle formed by two segments of ACPs (*θ*_0,ACP_ = 0 rad) and the equilibrium length of each ACP arms (*r*_0,ACP_ = 23.5 nm) are maintained by extensional (*κ*_s,ACP_) and bending (*κ*_b,ACP_) stiffnesses of ACPs, respectively.

An equilibrium angle formed by adjacent backbone segments (*θ*_0,M_ = 0 rad) and the equilibrium length of each motor backbone segment (*r*_s,M1_ = 42 nm) are maintained by bending (*κ*_b,M_) and extensional (*κ*_s,M1_) stiffnesses, respectively. The value of *κ*_s,M1_ is equal to that of *κ*_s,A_, whereas the value of *κ*_b,M_ is much larger than that of *κ*_b,A_ so that the backbone does not bend much. The extension of each motor arm is regulated by the two-spring model with the stiffnesses of transverse (*κ*_s,M2_) and longitudinal (*κ*_s,M3_) springs. The transverse spring maintains an equilibrium distance (*r*_0,M2_ = 13.5 nm) between the endpoint of a motor backbone where the motor arm is connected and an actin segment where the other end of the motor arm is bound, whereas the longitudinal spring maintains a right angle between the motor arm and the actin segment (*r*_0,M3_ = 0 nm).

Repulsive forces originate from the following harmonic potential (5):

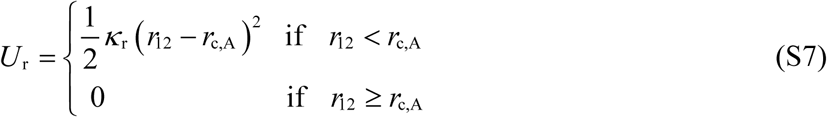

where *κ*_r_ is the strength of volume-exclusion effects, and *r*_12_ is a minimum distance between two neighboring actin segments. The repulsive force acts only if *r*_12_ is smaller than the diameter of actin segments (*r*_c,A_ = 7 nm).

Forces acting on actin segments due to bound ACPs and motors or due to the repulsive forces are distributed onto the two endpoints of the actin segments as described in our previous work in detail (6); as the endpoint is closer to the point of force application, it feels a larger fraction of the force.

### Dynamic behaviors of ACPs

The end of ACP segments binds to binding sites located on actin segments every 7 nm at a constant rate (*k*_+,ACP_) without preference for cross-linking angle. ACPs can unbind from actin filaments in a force-dependent manner, following Bell’s equation (7):

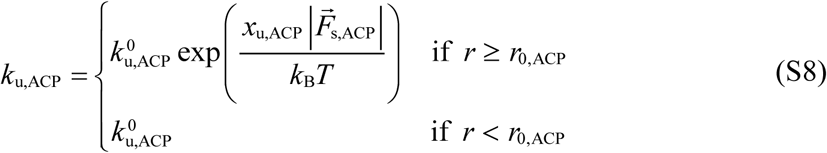

where 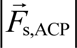 is the magnitude of a tensile spring force exerted on ACP segments, 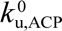 is the zero-force unbinding rate constant, and *x*u,ACP is sensitivity to applied force.

### Dynamic behaviors of motors

Motor arms bind to binding sites on actin segments at the rate of *k*_+,M_ = 40*N*_h_ s^-1^. The walking (*k*_w,M_) and unbinding (*k*_u,M_) rates of the motor arms are defined by the parallel cluster model (PCM) to account for the mechanochemical cycle of myosin II (9, 10). The PCM assumes three different states of myosin and 5 transition rates between the three states. One of them is a transition rate from the post-power-stroke state to the unbound state (*k*_20_), which requires the binding of ATP to myosin. The details of implementation and benchmarking of the PCM in our agent-based model are explained in our previous study (11). *k*_w,M_ and *k*_u,M_ calculated by the PCM tend to be smaller if a larger force is exerted on the longitudinal spring, meaning that the motor arms exhibit catch-bond behaviors. The unloaded (i.e., zero force) walking velocity and stall force (*F*_s_) of the motor arms with the reference value of *k*_20_ are ∼56 nm/s and ∼6*N*_h_ pN (= 24 pN), respectively. In some of the simulations, we varied *k*_20_ to change *k*_w,M_. As a result, *k*_u,M_ was also changed. In general, with higher *k*_20_, *k*_w,M_ and *k*_u,M_ increase (Fig. S6).

Unlike our previous studies, we additionally assumed that the motor arms immediately unbind from actin filaments if they feel a force greater than 30 pN. This assumption is necessary because the large stretch of the bundle (50% strain) is not possible if connections between motors and actin filaments do not break due to the catch-bond assumption. In fact, it has been reported that myosins exhibit a catch-slip bond behavior rather than a pure catch-bond behavior [62].

---

**Table S1.** List of all parameters used in our agent-based model. References are provided for some of the parameters if the parameter values were determined based on previous studies. “*” indicates reference values.

| Symbol | Definition | Value |
| --- | --- | --- |
| $r_{0,A}$ | Length of actin segments | $1.4 \times 10^{-7}$ [m] |
| $r_{c,A}$ | Diameter of actin segments | $7.0 \times 10^{-9}$ [m] (15) |
| $\theta_{0,A}$ | Bending angle formed by adjacent actin segments | 0 [rad] |
| $\kappa_{s,A}$ | Extensional stiffness of actin filaments | $1.69 \times 10^{-2}$ [N/m] |
| $\kappa_{b,A}$ | Bending stiffness of actin filaments | $2.64 \times 10^{-19}$ [N·m]<sup>*</sup> (4) |
| $r_{0,ACP}$ | Length of ACP segments | $2.35 \times 10^{-8}$ [m] (16) |
| $r_{c,ACP}$ | Diameter of ACP segments | $1.0 \times 10^{-8}$ [m] |
| $\theta_{0,ACP}$ | Bending angle formed by two ACP segments | 0 [rad] |
| $\kappa_{s,ACP}$ | Extensional stiffness of ACPs | $2.0 \times 10^{-3}$ [N/m] |
| $\kappa_{b,ACP}$ | Bending stiffness of ACPs | $1.04 \times 10^{-19}$ [N·m] |
| $r_{0,M1}$ | Length of motor backbone segments | $4.2 \times 10^{-8}$ [m] |
| $r_{0,M2}$ | Equilibrium length of the transverse spring for motor arms | $1.35 \times 10^{-8}$ [m] |
| $r_{0,M3}$ | Equilibrium length of the longitudinal spring for motor arms | 0 [m] |
| $r_{c,M}$ | Diameter of motor arms | $1.0 \times 10^{-8}$ [m] |
| $\theta_{0,M}$ | Bending angle formed by motor backbone segments | 0 [rad] |
| $\kappa_{s,M1}$ | Extensional stiffness of motor backbones | $1.69 \times 10^{-2}$ [N/m] |
| $\kappa_{s,M2}$ | Extensional stiffness 1 of motor arms (transverse) | $1.0 \times 10^{-3}$ [N/m] |
| $\kappa_{s,M3}$ | Extensional stiffness 2 of motor arms (longitudinal) | $1.0 \times 10^{-3}$ [N/m] |
| $\kappa_{b,M}$ | Bending stiffness of a motor backbone | $5.07 \times 10^{-18}$ [N·m] |
| $\kappa_r$ | Strength of repulsive forces | $1.69 \times 10^{-3}$ [N/m] |
| $N_h$ | Number of heads represented by a single motor arm | 4 |
| $N_a$ | Number of arms per motor | 8 |
| $k_{20}$ | ATP-dependent unbinding rate of myosin (for PCM) | $8$ [s<sup>-1</sup>]<sup>*</sup> |
| $k_{+,M}$ | Binding rate of motor arms | $40N_h$ [s<sup>-1</sup>] |
| $k_{+,ACP}$ | Binding rate of ACPs | $100$ [1/μM·s] |
| $k_{u,ACP}^0$ | Zero-force unbinding rate constant of ACPs | $0.05$ [s<sup>-1</sup>] |
| $x_{u,ACP}$ | Sensitivity of ACP unbinding to an applied force | $9.0 \times 10^{-10}$ [m] |
| $\kappa_{s,bnd}$ | Extensional stiffness for connection between filaments and boundaries | $1.0 \times 10^{-2}$ [N/m] |
| $\Delta t$ | Time step | $1.15 \times 10^{-6}$ [s] |
| $\mu$ | Viscosity of the surrounding medium | $8.6 \times 10^{-2}$ [kg/m·s] |
| $k_B T$ | Thermal energy | $4.142 \times 10^{-21}$ [J] |
| $L_f$ | Length of actin filaments | $4.9$ [μm] |
| $R_M$ | Ratio of motor concentration to $C_A$ | $0.01$<sup>*</sup> |
| $R_{ACP}$ | Ratio of ACP concentration to $C_A$ | 0.1 |
| $N_F$ | Total number of actin filaments | 50 |
| $\dot{\epsilon}_s$ | Stretching strain rate | 0.1 |
| $\dot{\epsilon}_c$ | Compressive strain rate | $-0.25$<sup>*</sup> |

## SUPPLEMENTAL FIGURES

**Fig. S1.**
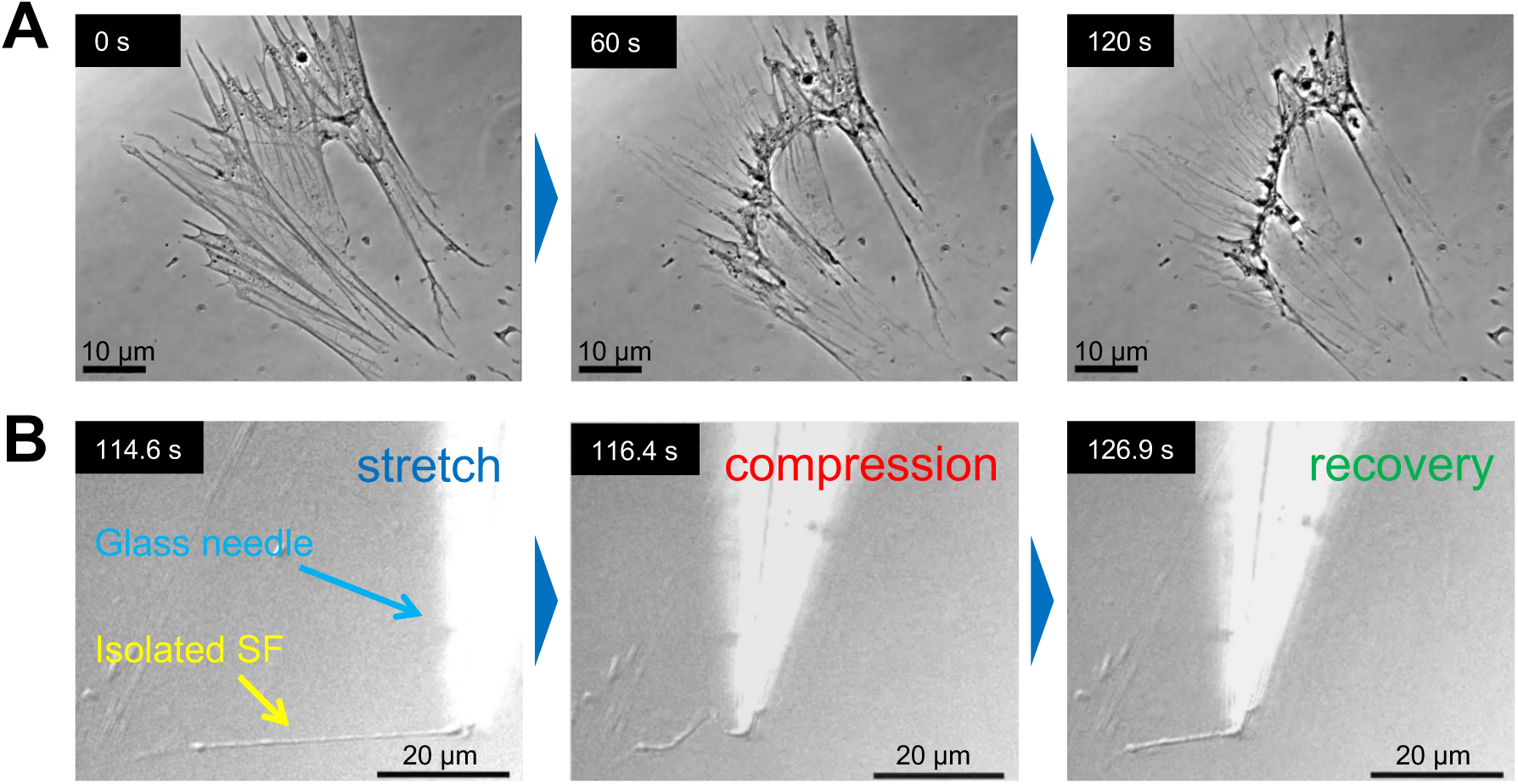
Mechanical behaviors of stress fibers (SFs) isolated from cells. (A) The contractility of isolated SFs could be maintained by adding the Mg-ATP buffer (1 mM) under the temperature of 25℃ and the ionic strength of 100 mM. (B) The buckling of isolated SFs was observed after the stretch-compression manipulation, which was followed by gradual recovery into a straight shape.

**Fig. S2.**
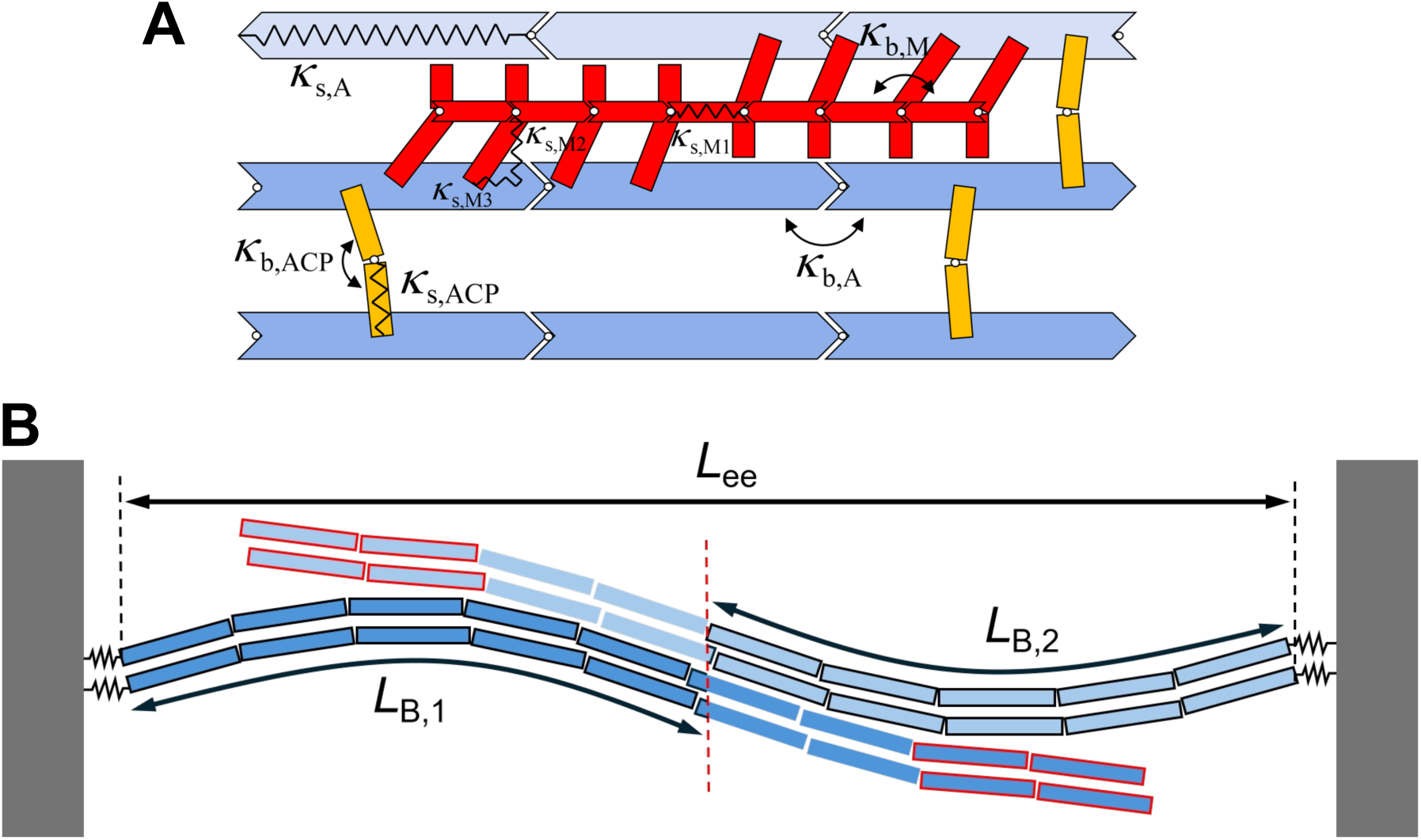
Agent-based model and quantification. (A) Actin filaments (blue), actin cross-linking proteins (ACPs, yellow), and motors (red) are simplified via cylindrical segments. Actin filaments are simplified into serially connected segments. ACPs consist of two segments connected at their center point. Motors have a backbone with 8 arms. ACPs transiently bind to actin filaments to connect pairs of actin filaments. Motor arms bind to actin filaments and walk toward to the barbed end of the filaments. *κ*_b_ and *κ*_s_ represent bending and extensional stiffnesses, respectively. Subscripts “A” and “M” represent actin and motor, respectively. (B) Schematic describing measurements, which is not drawn in real length-scale. The end-to-end distance of the bundle (*L*_ee_) is calculated by measuring a distance between the left-most and right-most endpoints of actin segments in the z direction. The contour length of the entire bundle (*L*_B_) is calculated by adding two contour lengths (*L*_B,1_ and *L*_B,2_) measured between each boundary and the midplane between two boundaries indicated by a red dashed line. Each of these two contour lengths is calculated by calculating the sum of the length of segments with black boundaries along each filament and then averaging the sums. *L*_B_ is calculated in this way to exclude the fraction of filaments separated from the main part of the bundle (with red boundaries near their pointed ends).

**Fig. S3.**
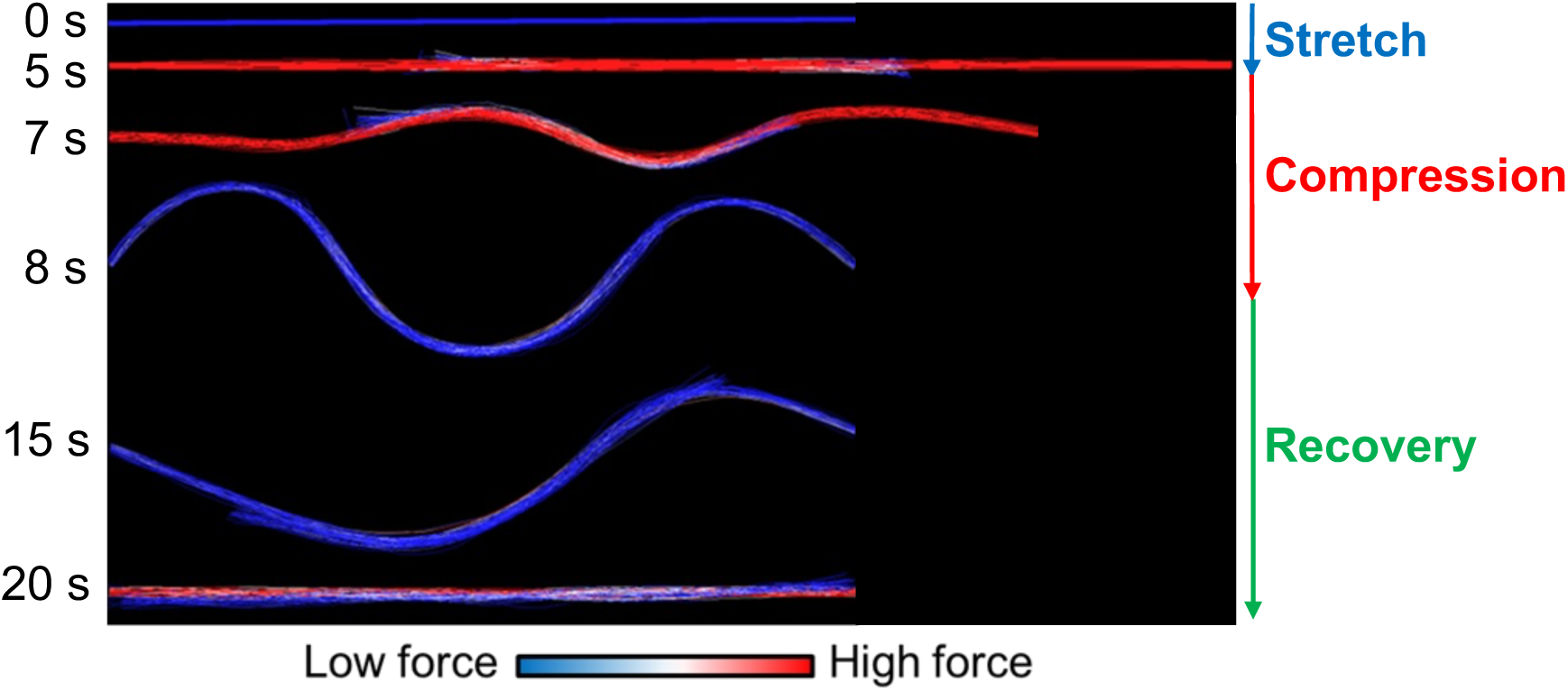
Level of a tensile force acting on the bundle in each phase visualized via the color scaling. A large tensile force is developed during the stretch phase. After the end of the compression phase, the tensile force is substantially smaller. The force is fully relaxed during the recovery phase. After full recovery into a straight shape, a tensile force is developed again due to motor activities.

**Fig. S4.**
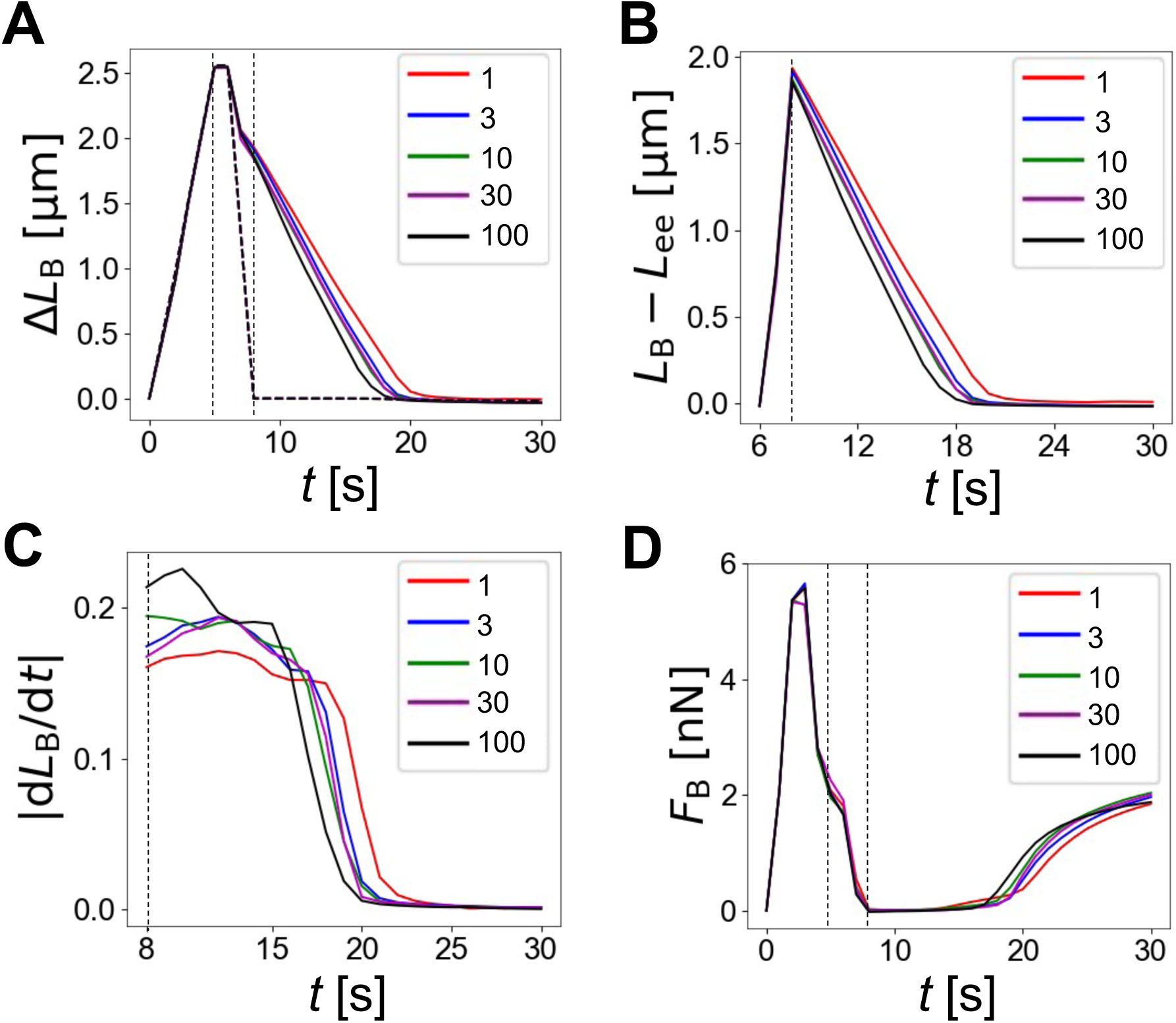
Analysis of cases with different filament bending stiffness (*κ*_b,A_). Numbers in the legends indicate filament bending stiffness relative to the reference one (*κ*_b,A_/*κ*^∗^). Vertical thin dashed lines represent the end of the stretch phase (at 5 s) and the end of compression phase (at 8 s). (A) A change in bundle contour length (Δ*L*_B_) over time. Thick dashed lines represent a change in the end-to-end distance (Δ*L*_ee_). (B) A difference between *L*_B_ and *L*_ee_ from the beginning of the compression phase. (C) The magnitude of the decreasing rate of *L*_B_ from the end of the compression phase. (D) Average tensile force acting on the bundle (*F*_B_) over time.

**Fig. S5.**
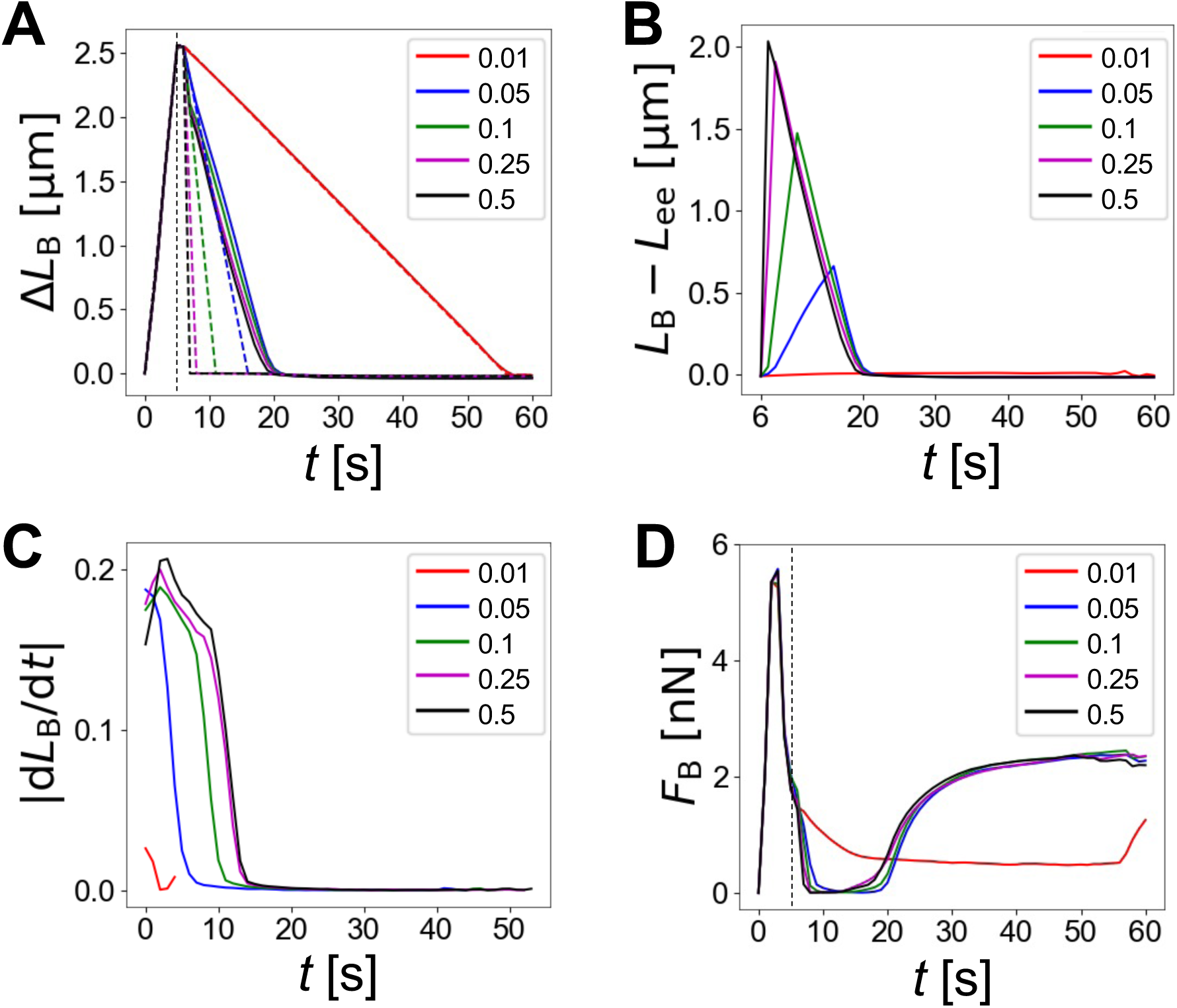
Quantification of cases with different compression rates (*ε̇*_c_). Numbers in the legends indicate the magnitude of *ε̇*_c_ in s^-1^. Vertical dashed lines represent the end of the stretch phase at 5 s. (A) A change in bundle contour length (Δ*L*_B_) over time. Thick dashed lines represent a change in the end-to-end distance (Δ*L*_ee_) in each case using the same color. (B) A difference between *L*_B_ and *L*_ee_ from the beginning of the compression phase. (C) The magnitude of the decreasing rate of *L*_B_ during the recovery phase. The starting time of the recovery phase for all cases is adjusted to 0 s for better comparison. (D) Average tensile force acting on the bundle (*F*_B_) over time.

**Fig. S6.**
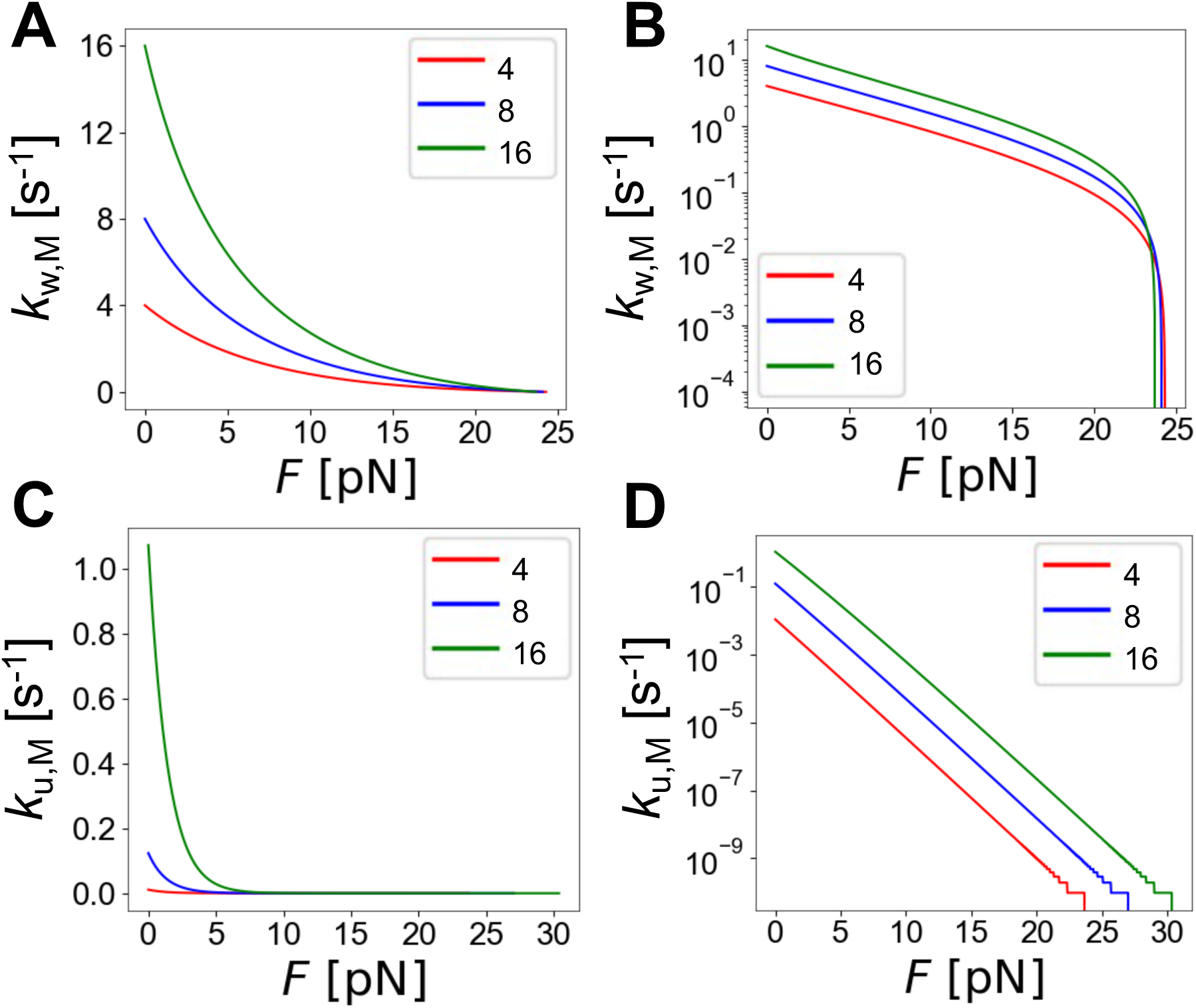
The force-dependent walking and unbinding rates of motors determined by the parallel cluster model with different values of the ATP-dependent unbinding rate (*k*_20_). (A, B) Walking rate with y axis in linear or log scale. The stall force corresponds to the x-intercept of each curve. (C, D) Unbinding rate with y axis in linear or log scale. With higher *k*_20_, the walking and unbinding rates are enhanced.

**Fig. S7.**
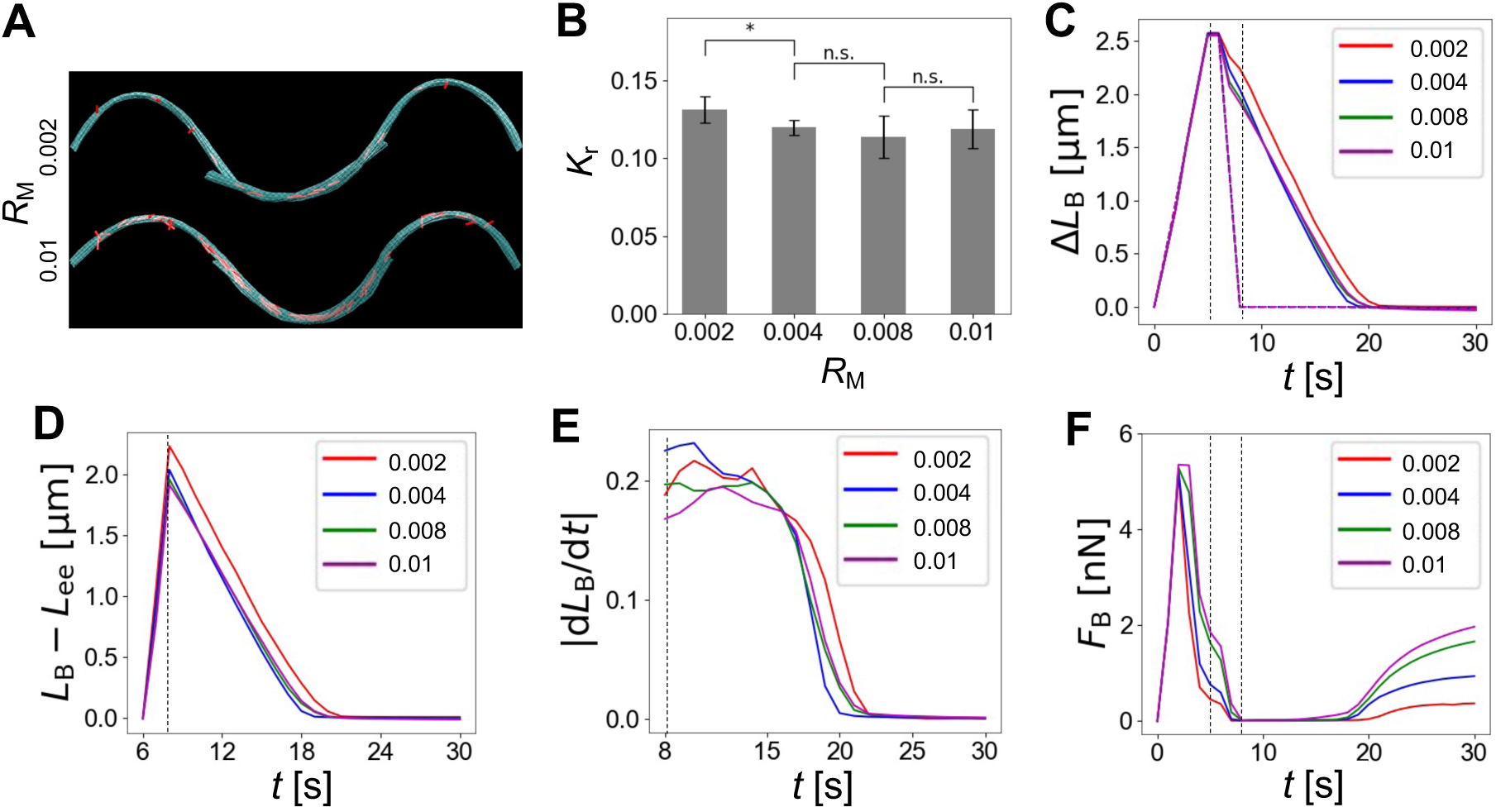
Analysis of cases with different values of the ATP-dependent unbinding rate for motors (*k*_20_). Numbers in the legends indicate the values of *k*_20_ in s^-1^. Vertical thin dashed lines represent the end of the stretch phase (at 5 s) and the end of compression phase (at 8 s). (A) The shape of the bundle at the end of the compression phase with two different *k*_20_. (B) Bundle curvature measured at the end of compression (*K*_r_). (C) A change in bundle contour length (Δ*L*_B_) over time. Thick dashed lines represent a change in the end-to-end distance (Δ *L*_ee_). (D) A difference between *L*_B_ and *L*_ee_ from the beginning of the compression phase. (E) The magnitude of the decreasing rate of *L*_B_ from the end of compression. (F) Average tensile force acting on the bundle (*F*_B_) over time.

**Fig. S8.**
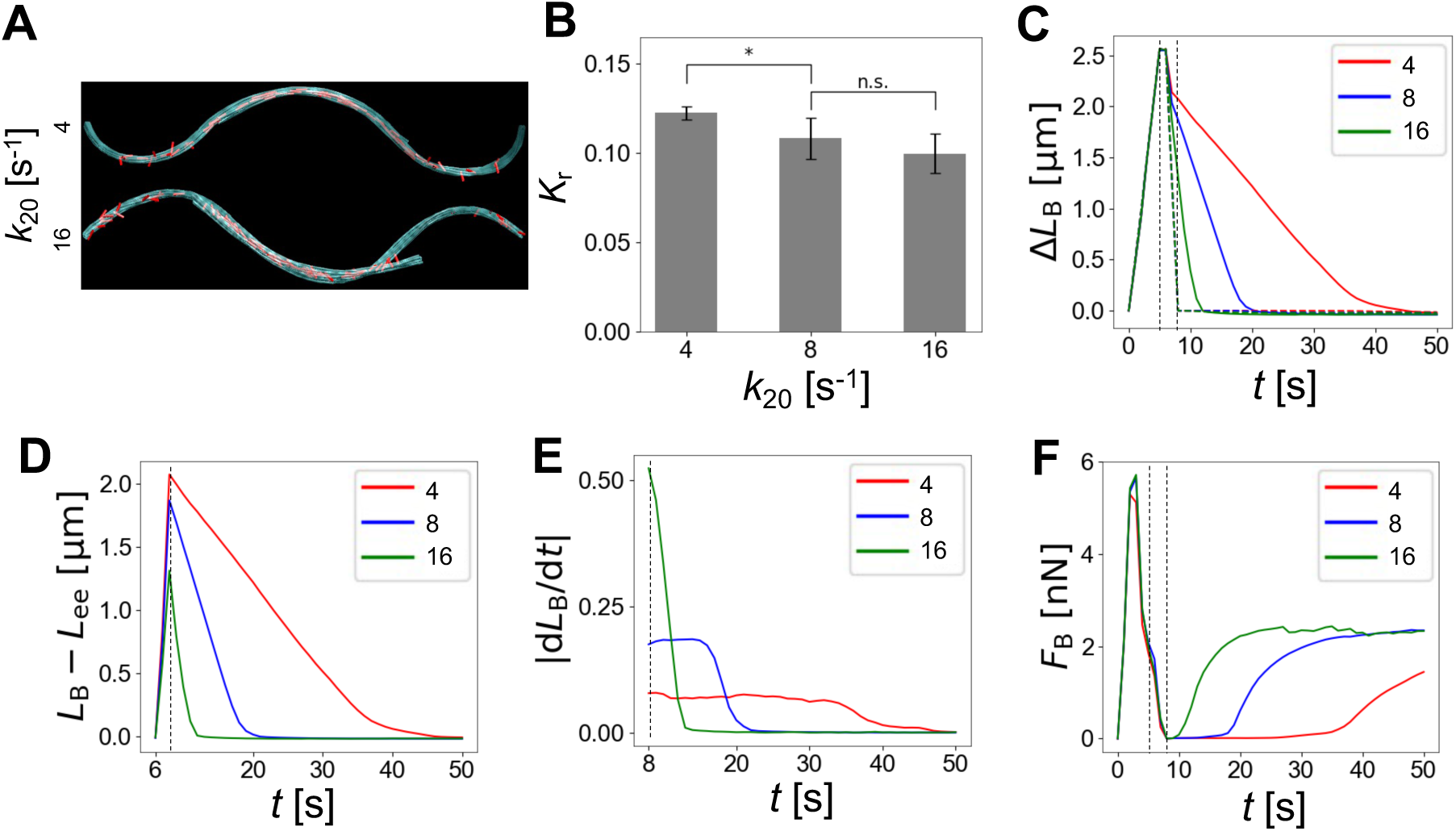
Quantification of cases with different motor density (*R*_M_). Numbers in the legends indicate the values of *R*_M_. Vertical thin dashed lines represent the end of the stretch phase (at 5 s) and the end of compression phase (at 8 s). (A) The shape of the bundle at the end of compression with two different *R*_M_. (B) Bundle curvature measured at the end of compression (*K*_r_). (C) A change in bundle contour length (Δ*L*_B_) over time. Thick dashed lines indicate a change in the end-to-end distance (Δ*L*_ee_). (D) A difference between *L*_B_ and *L*_ee_ from the beginning of the compression phase. (E) The magnitude of the decreasing rate of *L*_B_ from the end of compression. (F) Average tensile force acting on the bundle (*F*_B_) over time.

**Movie S1. Manipulation of individual SFs by a functionalized glass needle.** Various buckling patterns can be observed by altering the strain rate of the stretch-compression manipulation. The isolated SFs can return to the straight shape with various recovery time, denoting its different mechano-adaptive behaviors.

